# A single-cell immune atlas of primary and secondary lymphoid organs in pigs

**DOI:** 10.1101/2025.09.11.675613

**Authors:** Jayne E. Wiarda, Muskan Kapoor, Sathesh K. Sivasankaran, Kristen A. Byrne, Crystal L. Loving, Christopher K. Tuggle

**Affiliations:** Virus and Prion Research Unit, National Animal Disease Center, Agricultural Research Service, United States Department of Agriculture, Ames, IA, USA; Department of Animal Science, Iowa State University, Ames, IA, USA; Bioinformatics and Computational Biology Graduate Program, Iowa State University, Ames, IA, USA; Genome Informatics Facility, Iowa State University, Ames, IA, USA; Food Safety and Enteric Pathogens Research Unit, National Animal Disease Center, Agricultural Research Service, United States Department of Agriculture, Ames, IA, USA

**Keywords:** FAANG_1_, scRNA-seq_2_, immune_3_, atlas_4_, pig_5_, lymphoid_6_, single-cell_7_, cell annotation_8_

## Abstract

Single-cell RNA sequencing (scRNA-seq) has revolutionized understandings of cellular identities and functions due to the ability to study transcriptome-wide gene expression within individual cells. Multi-tissue scRNA-seq atlases have generated holistic understandings of body-wide cell dynamics and serve as key foundational resources for further scientific studies across a variety of species. Pigs are a valuable biomedical model, and pork is an essential global food source, but minimal understanding of immune cell identities and functions across anatomical locations limits agricultural and health advancements in pigs. To address current limitations, we apply scRNA-seq to create an atlas of immune cells recovered from key immune tissues including primary lymphoid organs (bone marrow and thymus) and secondary lymphoid organs (lymph node and spleen). Thymus data was compared to a previously published scRNA-seq dataset of pig thymus and shared a general consensus while also identifying several new thymic cell populations. Comparison of spleen to a human splenic scRNA-seq dataset also revealed conserved features, including two subsets of innate lymphoid cells conserved between pigs and humans. Spatial reconstruction of lymph node structure from scRNA-seq data revealed follicular organization with similar cell type distributions and cell signaling interactions to those in human lymph nodes. To expand accessibility of the scRNA-seq atlas for biological query, we deploy an interactive application and demonstrate its use for non-computational exploration of diverse cell populations recovered from bone marrow. Overall, results expand current foundational understandings of immune cell identities and functions in pig lymphoid organs and demonstrate pig-to-human immune similarities to consider for future research applications. Materials associated with this work are made readily accessible for others to investigate individual queries requiring foundational knowledge pertaining to pig immunity.

## 1 Introduction

The immune system is a complex orchestration of coordinating cells, vessels, and molecules that coalesce to prevent or control damages caused by pathogens, toxins, and other harmful insults. Lymphoid organs are the central hubs providing strategic localizations and specialized microenvironments needed to bolster immunity through production and activation of immune cells and immune mediators. Primary lymphoid organs serve as sites for immune cell production and development, while secondary lymphoid organs are sites where immune reactions are initiated and deployed. However, different primary and secondary lymphoid organs serve unique functions that are supported by distinct cell types [1]. As cells are the building blocks of immune reactions, an understanding of cellular heterogeneity across primary and secondary lymphoid organs is key to gaining a holistic interpretation of immune cell function and coordination. In this work, pigs were used to study cellular heterogeneity across lymphoid organs due to their agricultural importance as a food source [2] and their suitability as a biomedical model [3, 4].

To best survey the plethora of immune cells found across primary and secondary lymphoid organs in pigs, a high-resolution, omics-level technology is required. Single-cell RNA sequencing (scRNA-seq) is a technology that can achieve a transcriptome-wide survey of gene expression within individual cells, thus revolutionizing the functional annotation of individual cells, cell types, and cell states across various organisms, tissues, organ systems, disease states, and a variated multitude of other biological systems and scenarios. scRNA-seq is a suitable technology for discovery-based cell analysis, as previous phenotypic knowledge of cells is not required, and protein-based immunoreagent availability is not a limiting factor [5, 6].

Annotation of cell types is crucial to accurately interpret scRNA-seq data. The annotation process requires both an accurately annotated genome for correct assignment of sequence data to expressed genes within each cell and cell-level transcriptomes from many different tissues, ages, and biological states. The porcine genome is sufficiently contiguous [7], and substantial bulk RNA-seq [8] and scRNA-seq transcriptomic resources are available and accumulating, respectively, to provide sufficient information for porcine cell type annotation. Wang and colleagues recently published a multi-tissue atlas of 11 adult pig tissues, with the brain subdivided into another nine components [9]. While these tissues contained diverse cell types, this study was limited in immune tissue coverage, with only spleen and peripheral blood mononuclear cells (PBMCs) represented. Another recent report by Chen et al. created a scRNA-seq atlas of seven pig tissues, including two types of fat and many organs overlapping with the work by Wang et al. [10]. However, this work again was not immunologically focused, with spleen being the only lymphoid organ analyzed. Another recent pig scRNA-seq atlas by Zhang et al. was immunologically focused (spleen, lymph node, Peyer’s patches), but work was performed in germ-free and specific pathogen-free miniature pigs that aren’t representative of conventional livestock, and no primary lymphoid organs were analyzed in the work [11]. Others have focused on important porcine immune tissues such as thymus [12], gut-associated organized lymphoid tissues (i.e. Peyer’s patches) [13–15], spleen [16], and PBMCs [5, 17, 18]. While multi-tissue atlases are extremely useful for understanding the full embodiment of swine cell biology, and current individual immune tissue surveys have lent higher-resolution immune understandings in pigs, an immune-focused, more comprehensive multi-tissue assessment of pig leukocytes is still lacking but needed to gain comprehensive insights into immune coordination across lymphoid organs.

In this work, we aim to address important gaps in understanding the porcine immune cell landscape by establishing an immune organ-centric, multi-tissue scRNA-seq atlas. Tissues from both primary and secondary lymphoid organs were utilized to understand immune cellular dynamics at primary lymphoid sites of immune cell production and development (bone marrow, thymus) and at secondary lymphoid effector sites where immune cells routinely function to activate versus regulate immune responses (spleen, lymph node). In this work, we establish functionally- and phenotypically-relevant annotations for immune cells identified in each lymphoid organ, demonstrate applicable ways each tissue dataset can be utilized to further resolve immunological functions in swine, and draw several similarities of our findings to human immune components, thus suggesting potential suitability of pigs for several areas of biomedical study.

## 2 Materials and methods

### 2.1 Animals and sample collection

Four-week-old Yorkshire piglets were delivered to the National Animal Disease Center (NADC) in Ames, Iowa and housed in a biosecurity level (BSL)-2 room. Pigs were initially given commercial starter feed and then transitioned to a commercial grower/finisher feed after approximately two weeks. At approximately six months of age, two non-castrated male pigs were euthanized with intravenous barbiturate (sodium pentobarbital) overdose via an auricle vein. Immediately postmortem, four rib bones and sections of spleen, thymus, and ileocecal lymph node were collected. Ribs were used to isolate bone marrow cells. Other tissues were collected for cell isolations and *in situ* staining as described in subsequent sections. Animal experiments were performed according to procedures approved by the Institutional Animal Care and Use Committee at the NADC.

### 2.2 Cell isolations and cryopreservation

Collected spleen, thymus, and ileocecal lymph node tissues (∼2 g) were transported back to the lab in 10 mL tissue buffer (2 mM EDTA [Invitrogen AM9260G], 2mM L-glutamine [Gibco 25-030], 0.5% bovine serum albumin [BSA; Sigma-Aldrich A9418] in Hank’s Balanced Salt Solution [HBSS; Gibco 14175]) in a gentleMACS C Tube (Miltenyi 130-093-237). In the lab, tissues were mechanically homogenized using a gentleMACS Octo Dissociator (Miltenyi 130-095-937) with the programed spleen – cells protocol. Cell suspensions were passed through a 100-micron nylon mesh cell strainer, and strainers were washed with 10 mL tissue buffer. Cells were pelleted by centrifugation at 300 xg for 5 min room temperature (RT). Cell pellets were resuspended in residual volume (<1 mL) and then incubated with 20 mL ACK lysis buffer (ThermoFisher A1049201) for 3 min to lyse red blood cells and centrifuged again at 300 xg for 5 min RT. Spleen cells were treated with ACK lysis buffer a second time. Cells were resuspended in 10 mL tissue buffer and passed through a 70-micron nylon mesh cell strainer. Cells were centrifuged 300 xg for 5 min RT and resuspended in tissue buffer.

Bone marrow cells were isolated by flushing ∼30 mL tissue buffer through two rib bones using an 18 gauge needle with 10 mL syringe and collecting cells into a 50 mL conical as flushed out. Recovered cells were pelleted by centrifuging 300 xg for 5 min RT. Cells were then passed through cell strainers and treated with ACK lysis buffer as described above.

Cell quantity and viability was assessed using the Muse Count & Viability Assay Kit (Luminex MCH100102) with a Muse Cell Analyzer (Luminex 0500-3115). To further enrich for live cells, cell suspensions were processed through a Dead Cell Removal Kit (Miltenyi 130-090-101) according to manufacturer’s protocol with a starting quantity of 5×10^7^ total cells as previously described [14]. Recovered cells were cryopreserved according to the 10X Genomics Sample Preparation Demonstrated Protocol in multiple aliquots of 1×10^7^ cells per vial.

### 2.3 Single-cell RNA library preparation sequencing

Cells were thawed according to the 10X Genomics Sample Preparation Demonstrated protocols, then processed twice (spleen, thymus, lymph node) or once (bone marrow) through the Dead Cell Removal Kit. Cell quantity and viability was assessed as described above and deemed adequate for scRNA-seq (>75% live cells per sample). Partitioning and library preparation were performed according to the Chromium Single Cell 3’ Reagent Kits v2/3 User Guide (10X Genomics CG00052), and 100 base paired-end sequencing was performed on a HiSeq3000 (Illumina) at the Iowa State University DNA Facility. Aliquots of bone marrow cells from both pigs and lymph node cells from one pig were thawed and processed for an additional scRNA-seq submission as described above. Library preparation reagents used v2 or v3 chemistry as follows: bone marrow samples and the lymph node sample processed for an additional submission used v2 chemistry in the first run and v3 chemistry in the second run; thymus cells used v3 chemistry; the lymph node sample processed only once used v3 chemistry; spleen samples used v2 chemistry. Data integration (described in subsequent methods) was used to alleviate effects of submission batch and library chemistry. Sequencing data were deposited as .fastq files for both forward and reverse strands following image analysis, basecalling, and demultiplexing.

### 2.4 Data analysis

#### 2.4.1 Initial data processing

Initial data processing was performed similar to previous work [5]. Briefly, reads were aligned to the *Sus scrofa* 11.1 reference genome and v97 annotation file [19] with Cell Ranger v4.0 (10X Genomics). Single-cell gene counts were next analyzed with SoupX v1.4.5 [20] to calculate and remove contaminating ambient RNA. Genes with sum-zero expression across all samples were removed from the dataset, as were cells with <=300 total genes, <=500 UMIs, and/or >= 10% mitochondrial reads detected. Scrublet v0.2 [21] was used to estimate doublet probabilities, and cells with doublet probabilities >=0.25 were removed. Filtered data were then further analyzed with Seurat v4.3.0.1 [22, 23] to perform data normalization (SCT and log transformations), integration, dimensionality reductions (PCA, t-SNE, UMAP), nearest neighbor calculations, and clustering as previously described [5], treating each tissue as an individual dataset. SCT-normalized data were used for data integration, and integrated data were used for PCA. The number of principal components used for additional dimensionality reductions (t-SNE, UMAP), finding nearest neighbors, and hierarchical clustering were calculated as previously described [5].

#### 2.4.2 Cell annotation

Clustering was performed in Seurat as previously described [5]. Clustering resolutions were tested in intervals of 0.5 for each tissue dataset until sufficient segregation of cell types was achieved for annotation. Cell clusters were annotated by assessing canonical gene expression and lists of differentially expressed genes (differential gene expression analysis described below). At times, multiple clusters were converged into a single cell type annotation due to overlapping transcriptional profiles used to describe cell types at a biological level. In bone marrow, a single cluster containing progenitors, B cells, and myeloid cells could not be segregated into distinct cell lineages by increasing clustering resolution (clustering resolution tested from 0.5 to 10 at 0.5 intervals), so a data subset of cells from the single cluster was created and re-processed as a new dataset (normalization, integration, dimensionality reduction, nearest neighbor calculation, clustering) to identify cell types, and cell identities for the single cluster were incorporated back into the original bone marrow dataset containing all cells. In bone marrow and spleen datasets, a cluster containing high levels of hemoglobin genes (*HBA\**, *HBB*) and conflicting expression profiles of lineage-specific markers were identified. Creating subsets of data containing cells with high hemoglobin expression and performing re-clustering did not further indicate cell identity, segregate conflicting lineage-specific markers, or suggest an alternative cell identity (e.g. progenitor cells). We therefore removed these cells from further analyses and propose these clusters may be aggregates of cells adhered to erythrocytes, enclosed in the same droplet as erythrocytes during cell partitioning, or another form of unknown technical artifact.

#### 2.4.3 Data merging of cell lineages

From each tissue dataset, data subsets were created for each of myeloid, B, and T/ILC lineage cells similar to previous work [14]. Cell lineage data subsets from each tissue were then merged into a new Seurat object to create new datasets containing all cells of each single lineage. Cells were processed through data normalization, integration, PCA calculations, and dimensionality reduction as described in methods for initial data processing.

#### 2.4.4 Reference-based cell type prediction and mapping

Reference-based cell type prediction and mapping was performed with defined query and reference datasets as described in previous works [14, 24, 25]. Thymus and spleen datasets were treated as query datasets and mapped separately to previously published datasets of porcine thymus [12] or human spleen [26], respectively, that were treated as reference datasets. For reference mapping to human data, pig single-cell data was humanized as previously described [14] prior to performing reference-based cell type prediction and mapping. A Seurat object of reference data for previously published porcine thymus was obtained by requesting data from the corresponding author [12]. A Seurat object of reference data for previously published human spleen was obtained as outlined in the article’s data availability statement [26]. Canonical correlation analysis (CCA) reduction methods were used to identify prediction/mapping anchors.

#### 2.4.5 Data merging of innate lymphoid cells from human and pig spleen

Cells annotated as cytotoxic ILCs or *NCR1*+*EOMES*+ ILCs in our humanized spleen dataset and *CD160*+ NK or *FCGR3A+* NK cells in the human spleen dataset [26] were extracted and merged into a single Seurat object. Cells were processed through data normalization, integration, PCA calculations, dimensionality reduction, and hierarchical clustering as described in methods for initial data processing.

#### 2.4.6 Cell-cell interaction network analysis

CellChat v2.1.2 [27] was used to identify cellular communication networks as previously described [13]. The human CellChat database was used to identify ligand- and receptor-encoding genes after porcine data from our lymph node dataset was humanized as previously described [14]. Only cell-cell interactions were used for analyses.

#### 2.4.7 Three-dimensional tissue organization reconstruction

CSOmapR v1.0 [28] was applied to the humanized lymph node tissue dataset used for cell-cell interaction analysis above. The human CellChat database was used to identify ligand- and receptor-encoding genes that were applied to create a 3D reconstruction of lymph node cell organization based on receptor and ligand expression profiles.

### 2.5 Shiny app development

The Shiny R package for immune tissue cell data visualization is a collection of components that facilitates the interactive representation and viewing of R analysis results [29]. The Shiny-PIGGI (https://shinypiggi.ansci.iastate.edu) is implemented completely in R, runs on any modern web browser, and requires no programming. Our main goal was to develop an interactive web application that allows users such as animal scientists and immunologists to visualize biological datasets. We utilized the R package Seurat and several other R packages (shiny, shiny dashboard, shiny themes, uwot, DT shinycssloaders, shinydisconnect, shiny alert, HTML tools, and HTML widgets) to design the user interactive interface. For differential expression, we used Wilcoxon rank sum test via the presto package [30]. Users can select any two cell types across tissues for further tissue-based cell type comparisons. The results are displayed in a table and can be downloaded based on filtering within the app. For gene expression visualization, we used feature and violin plot to visualize expression of a query gene across 4 tissues. Next, Seurat based reference mapping was implemented, using bone marrow as a reference tissue, and the other three tissues serves as query datasets. For each predicted cell type, mapping scores, prediction scores, and predicted cell annotations were computed and visualized. Additionally, we provided a link to downloadable .cloupe and .h5seurat files converted from the Seurat objects and available through Ag Data Commons. .cloupe files are compatible with interactive data query features available from Loupe Cell Browser (10X Genomics) to facilitate easy visualization/analyses outside shiny app. .h5seurat files maintain all features of Seurat objects used for data analysis herein and are compatible with upload into R environments using SeuratDisk but are also convertible to data object formats compatible with other computing environments via additional file conversions.

### 2.6 *In situ* staining

Sections of lymph node tissues were cut to appropriate size, placed into a cassette, fixed in 10% neutral-buffered formalin (3.7% formaldehyde) for ∼48 h RT, transferred to 70% ethanol, and embedded in paraffin blocks within a week of collection. Embedded tissues were cut into four-micron thick sections and adhered to Superfrost Plus charged microscope slides (Fisherbrand 12-550-15). RNA *in situ* hybridization to detect T receptor delta constant (*TRDC*) transcript was performed using a Sus scrofa *TRDC* probe (Advanced Cell Diagnostics 553141) and RNAscope 2.5 HD Detection Kit (Advanced Cell Diagnostics 322360) as previously described [31]. Immunohistochemistry to detect CD3ε protein was performed using a polyclonal rabbit anti-human CD3ε antibody (Dako A0452) as previously described [32]. Serial sections were used for *TRDC* and CD3ε labeling of lymph node tissue.

## 3 Results

### 3.1 An immune atlas of lymphoid organs in pigs

Single-cell RNA sequencing (scRNA-seq) was performed using cell fractions isolated from primary lymphoid organs (bone marrow, thymus) and secondary lymphoid organs (spleen, ileocecal lymph node) of two ∼6-month-old intact male Yorkshire pigs. Following quality control processing, final single-cell datasets included 5,899 bone marrow-derived cells (**Figure 1A**), 17,940 thymus-derived cells (**Figure 1B**), 20,210 lymph node-derived cells (**Figure 1C**), and 5,621 spleen-derived cells (**Figure 1D**), totaling overall at 49,670 cells that were further analyzed and annotated. Canonical gene expression patterns were used to annotate cell types (**Figure 1E-H**, **Figure 2A-C**). Progenitor cells were identified in primary lymphoid organs by *KIT* expression [33] and lack of lineage commitment genes (**Figure 1E-F**), with the majority identified in bone marrow (**Figure 1A**). Lineage-committed and/or developmentally mature immune cells belonging to T/innate lymphoid cell (ILC), B/antibody-secreting cell (ASC), and myeloid lineages were also identified in all tissues by expression of lineage-specific/lineage-enriched genes (*CD3E, CD3G, CD3D, CD247, ZAP70* for T/ILC; *CD79B, CD19, PAX5, JCHAIN* for B/ASC; *AIF1, CST3* for myeloid [5, 14] (**Figure 1E-H**, **Figure 2A-C**). Except for myeloid lineage-derived osteoclasts in the bone marrow (**Figure 1A**), stromal and epithelial cells were not identified, likely due to cell isolation methods catering to leukocyte rather than stromal/epithelial cell isolation (lack of enzymatic tissue digestion [34]) and cryopreservation protocols [35]). Overall, scRNA-seq analysis revealed both conserved and distinct transcriptional features of leukocytes derived from primary lymphoid organs (bone marrow, thymus) and secondary lymphoid organs (lymph node, spleen) of pigs.

**Figure 1.**
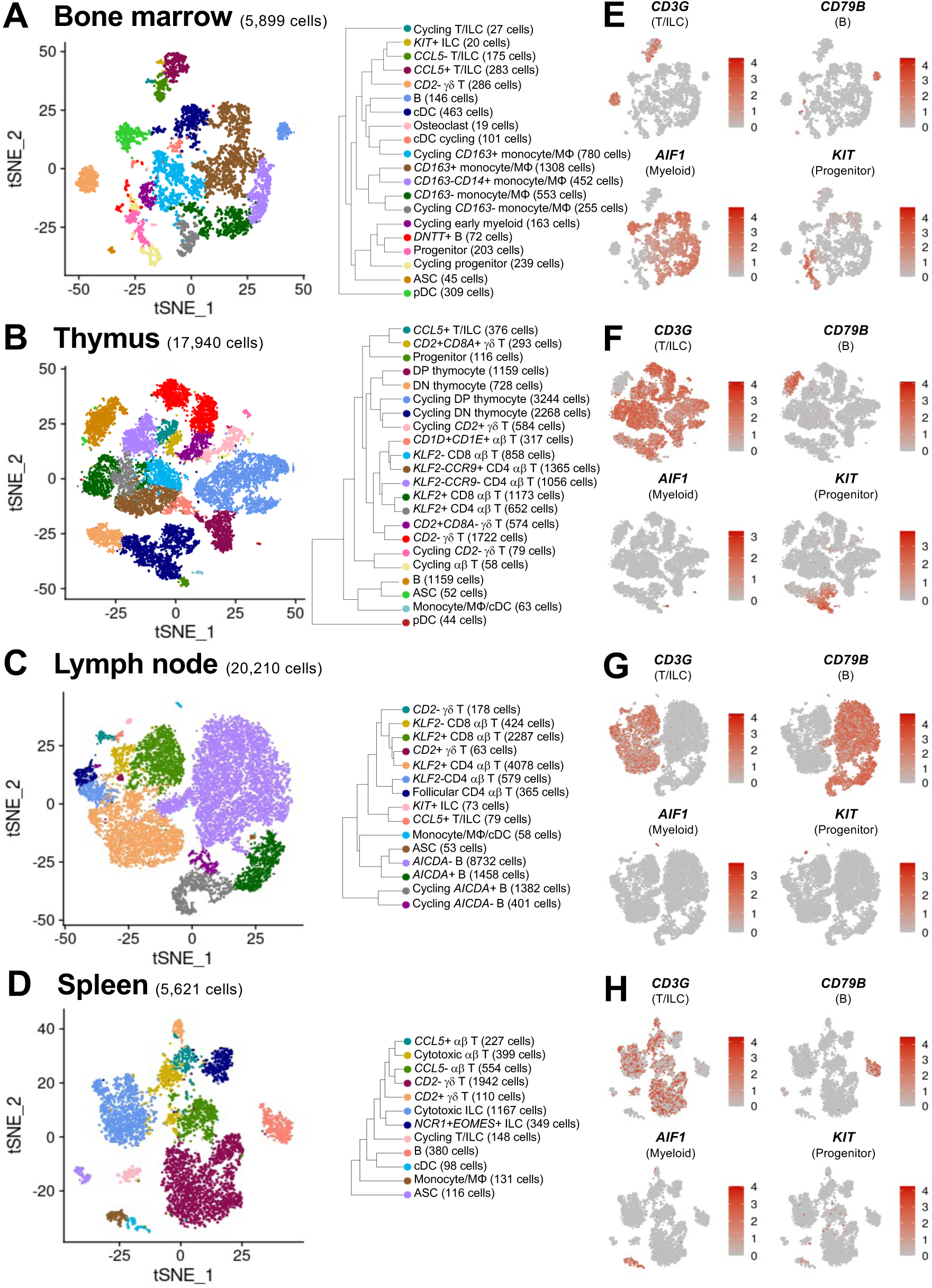
Cell types identified in porcine primary and secondary lymphoid tissues via scRNA-seq. (A-D) t-SNE plots of cell types identified in scRNA-seq datasets of porcine bone marrow (A), thymus (B), lymph node (C), and spleen (D). Each individual point represents one cell, and the color of each point corresponds to annotated cell identity. On the right of each t-SNE plot is a hierarchical tree indicating relatedness of annotated cell types in each dataset. (E-H) t-SNE plots of expression levels for selected canonical genes in scRNA-seq datasets of porcine bone marrow (A), thymus (B), lymph node (C), and spleen (D). Each individual point represents one cell, and the color of each point corresponds to relative expression level of the indicated gene. Abbreviations: ASC (antibody-secreting cell); cDC (conventional dendritic cell); DN (double negative); DP (double positive); ILC (innate lymphoid cell); MΦ (macrophage); pDC (plasmacytoid denritic cell); scRNA-seq (single-cell RNA sequencing); t-SNE (t-distributed stochastic neighbor embedding)

**Figure 2.**
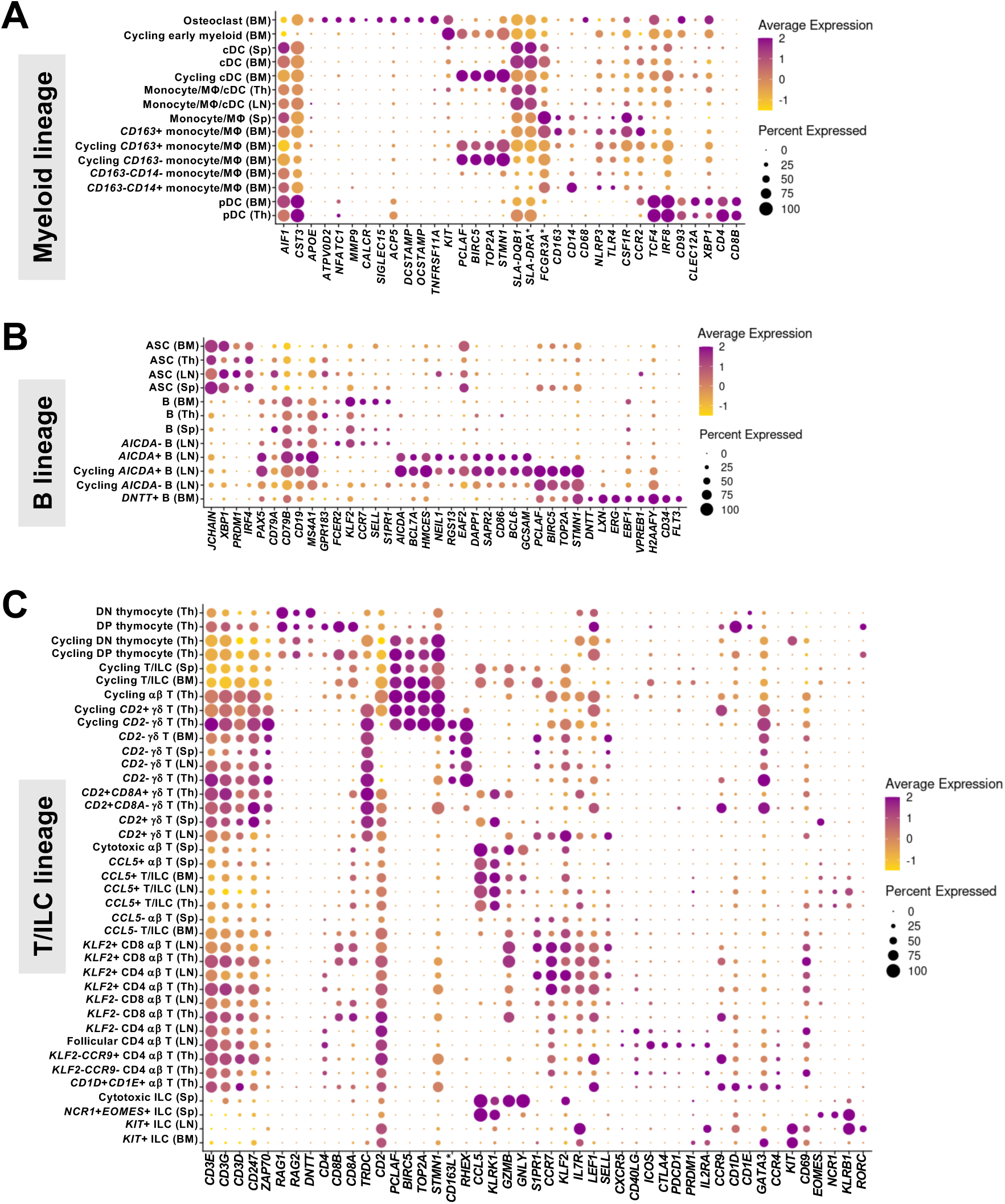
Canonical gene expression in myeloid lineage, B lineage, and T/ILC lineage cells recovered from porcine primary and secondary lymphoid tissues via scRNA-seq. (A-C) Dot plots of canonical gene expression (x-axes) in annotated cell types (y-axes) of myeloid lineage (A), B lineage (B), and T/ILC lineage (C) cells recovered from porcine bone marrow, thymus, lymph node, or spleen (cells shown in Figure 1A-D). Dot size indicates the percentage of an annotated cell type expressing a gene. Dot color indicates average gene expression level for those cells expressing a gene within an annotated cell type relative to all other cells in the same lineage. Abbreviations: ASC (antibody-secreting cell); BM (bone marrow); cDC (conventional dendritic cell); DN (double negative); DP (double positive); ILC (innate lymphoid cell); LN (lymph node); MΦ (macrophage); pDC (plasmacytoid denritic cell); scRNA-seq (single-cell RNA sequencing); Sp (spleen); Th (thymus)

#### 3.1.1 Myeloid lineage leukocytes

The majority of all recovered myeloid lineage leukocytes were derived from bone marrow (**Figure 1A**), and smaller populations were also identified in thymus, spleen, and lymph node (**Figure 1B-D**). Canonical gene expression used to identify myeloid lineage cells is shown in **Figure 2A** and discussed below. Though pDCs are derived from lymphoid progenitors [36], these cells also share transcriptional and functional commonalities with myeloid cells and are included in **Figure 2A**.

*AIF1* and *CST3* were used as myeloid lineage identifiers constitutively expressed across all pDC, cDC, and monocyte/macrophage (monocyte/MΦ) populations with the exception of a population of osteoclasts (lacking *AIF1*) and cycling early myeloid cells (lacking *AIF1* and *CST3* expression), both located in bone marrow. Osteoclasts notably expressed several highly specific and previously-reported genes, including *ATP6V0D2, NFATC1, MMP9, CALCR, SIGLEC15, ACP5, DCSTAMP, OCSTAMP, TNFRSF11A* [37–39]. Strong expression of *KIT* indicated cycling early myeloid cells were recently committed to the myeloid lineage, likely associated with hematopoietic development in the bone marrow [33, 40, 41]. Cycling early myeloid cells also had strong expression of cell cycle markers, such as *PCLAF, BIRC5, TOP2A, STMN1* [42, 43], indicating further that they were a cycling cell population.

cDCs were identified in all tissues by expression of *FLT3, FCER1A,* and high relative expression of MHC-II-encoding genes (*SLA-DQB1, SLA-DRA\**) [5, 14]. The only cycling population of cDCs was identified in bone marrow and expressed cell cycle genes such as *PCLAF, BIRC5, TOP2A, STMN1*. In thymus and lymph node, cDC and monocyte/MΦ populations were small and therefore combined into a single annotation due to limitations of clustering granularity.

Monocyte/MΦ populations had strong expression of *LYZ* and mixed expression of *FCGR3A*, CD163, CD14, CD68, NLRP3, TLR4, CSF1R, CCR2*. The majority of monocyte/MΦ populations were identified in bone marrow, where all populations constitutively expressed *FCGR3A\** (encoding CD16) but had variable expression of *CD163* and *CD14* that was used to discriminate populations further. Use of *CD163* as a marker for discrimination of bone marrow monocyte/MΦ populations is similar to previous work, where *CD163* expression was used as a discriminate between resident and non-resident bone marrow MΦs [44]. Expression of cell cycle genes (*PCLAF, BIRC5, TOP2A, STMN1*) was used to further identify cycling *CD163*+ and *CD163*-bone marrow monocyte/MΦ cells.

pDCs were identified by expression of *TCF4, IRF8, CD93, CLEC12A, XBP1, CD4, CD8B* [5, 14] in only thymus and bone marrow. Similar to monocyte/MΦ and cDCs, pDCs also expressed *AIF1* and *CST3*.

#### 3.1.2 B cells and antibody-secreting cells

ASCs expressing *JCHAIN, XBP1, PRDM1, IRF4* and B cells expressing *PAX5, CD79A, CD79B, CD19, MS4A1* [5, 14] were recovered in all tissues (**Figure 1A-D**), with the largest and most diverse B cell populations being found in lymph node (**Figure 1C**). Canonical gene expression used to identify B/ASC lineage cells is shown in **Figure 2B** and further discussed below.

Within lymph node, the majority of B cells were *AICDA*-B cells expressing *GPR183, FCER2, KLF2,* and circulatory genes *CCR7, SELL, S1PR1*, indicative of a resting cell state [13, 14]. Activation-associated genes (*AICDA, BCL7A, HMCES, NEIL1, RGS13, EAF2, DAPP1, S1PR2, CD86*) were indicative of somatic hypermutation, class switching, localization to germinal centers, and apoptosis [13, 45–49] and occurred in lymph node *AICDA*+ B cell populations. Expression of germinal center-associated genes (*BCL6, GCSAM*) [13, 50, 51] further suggested *AICDA*+ cells likely originated from active germinal centers. Cycling *AICDA*+ B cells were further annotated by expression of cell cycle genes (*PCLAF, BIRC5, TOP2A, STMN1*) that indicated probable locations within germinal center light and dark zones associated with stages of B cell activation and cycling [13]. An additional population of cycling *AICDA*-B cells lacking expression of many activation- and germinal center-associated genes was also identified in lymph node.

In bone marrow, a unique population of pro-/pre-B cells was also recovered, termed *DNTT*+ B cells. *DNTT+* B cells expressed genes associated with B cell lineage commitment and development (*DNTT, LXN, ERG, EBF1, VPREB1, H2AFY, CD34, FLT3*), indicating this was a population of recently committed B lineage cells undergoing development in the bone marrow, the primary site of B cell development [45, 52–55].

Remaining B cell populations in bone marrow, thymus, and spleen lacked expression of genes indicating B cell activation, germinal center localization, or development but did express genes in common with lymph node *AICDA*-B cells presumed to be in a resting state (*GPR183, CCR7, KLF2, SELL, FCER2, S1PR1*).

ASCs were also identified across all four tissues, and ASCs in bone marrow and spleen had higher expression of cell cycle genes (*PCLAF, BIRC5, TOP2A, STMN1*), suggesting a higher occurrence of plasmablasts.

#### 3.1.3 Thymocytes, T cells, and innate lymphoid cells

T cells expressed pan-T cell marker *CD3E* and other T cell receptor (TCR)-related genes (*CD3D, CD3G, CD247, ZAP70*), while ILCs were transcriptionally similar to T cells but lacked expression of *CD3E* [5, 14]. T cells and ILCs were identified in all analyzed tissues (**Figure 1A-D**). Canonical gene expression used to identify T/ILC lineage cells is shown in **Figure 2C** and discussed below.

Thymocytes were unsurprisingly unique to the thymus, the primary site of T cell development. All thymocyte populations expressed *RAG1, RAG2,* and *DNTT*, indicative of TCR rearrangement occurring during thymic T cell development [12, 56]. Double-positive (DP) thymocytes expressed *CD4*, *CD8A,* and *CD8B*, indicative of both CD4 and CD8 co-receptor expression, while double-negative (DN) thymocytes lacked such expression [12, 56]. Variable expression of *TRDC*, encoding for a portion of the TCR expressed by γδ T cells, was also noted in both DN and DP thymocyte populations. DP and DN thymocytes were further divided into cycling and non-cycling populations by expression of cell cycle genes (*PCLAF, BIRC5, TOP2A, STMN1*), leaving a total of four thymocyte populations identified.

Mature αβ T cells, γδ T cells, and ILCs were found in all tissues and lacked expression of genes associated with TCR rearrangement (*RAG1, RAG2, DNTT*). Clusters of cycling T cells and/or ILCs were identified in all samples except lymph node by expression of genes associated with cellular replication and division (*PCLAF, BIRC5, TOP2A, STMN1*). In bone marrow and spleen, cycling T cells and ILCs were grouped into single populations, while in thymus, where the largest number of total T/ILC populations were defined, cycling T cells were divided into αβ T cells and two subsets of cycling γδ T cells discussed subsequently.

γδ T cells (*TRDC*+) were separated into *CD2*+ and *CD2*-subsets in all tissues, mirroring conventional porcine γδ T cell classifications used in previous porcine-specific scRNA-seq work [5, 14]. *CD2*-γδ T cells were found in every tissue, and five total *CD2*-γδ T cell populations were annotated, including a population of cycling *CD2*-γδ T cells in thymus and populations of non-cycling *CD2*-γδ T cells in each of thymus, bone marrow, lymph node, and spleen. Similar to previous works [5, 14], porcine *CD2*-γδ T cells had strong expression of genes *CD163L\** and *RHEX* compared to other T cells across all tissues, suggesting these genes as a conserved expression signature for *CD2*-γδ T cells throughout anatomical locations of the pig.

Five populations of *CD2*+ γδ T cells were also identified, including three populations in thymus and single populations in lymph node and spleen. In thymus, two populations were comprised of non-cycling *CD2+* γδ T cells that were further divided into *CD8A+* and *CD8A-* populations, corresponding to conventional porcine γδ T cell classifications also used in previous porcine-specific scRNA-seq work [5]. Thymic CD2+ CD8A+ γδ T cells had increased expression of cell activation and/or effector-associated genes previously reported in PBMCs [5], including *KLRK1, CCL5,* and *GZMB*. Cycling *CD2*+ γδ T cells in thymus were also *CD8A*- and lacked expression of *KLRK1, CCL5,* and *GZMB*. *CD2*+ γδ T cell populations found outside of the thymus included splenic *CD2*+ γδ T cells that had strong expression of activation/effector genes *KLRK1, CCL5,* and *GZMB* and lymph node *CD2*+ γδ T cells that lacked such expression.

*CCL5*, an inferred marker of effector cell status [14, 57], was used to further discriminate populations of non-cycling αβ T cells (lacking *TRDC* expression) and ILCs (lacking *CD3E* and other TCR-related gene expression) in all tissues as *CCL5*+ or *CCL5*-populations. In bone marrow, αβ T cells and ILCs were included in the same populations due to the low number of total αβ T cells and ILCs identified and included clusters of *CCL5*+ T/ILCs and *CCL5*-T/ILCs in addition to the cycling T/ILC cluster and γδ T cell clusters mentioned above. In spleen, two populations of non-cycling αβ T cells were *CCL5*+ and further divided by expression (cytotoxic αβ T cells) or absence in expression (*CCL5*+ αβ T cells) of cytotoxicity-related genes (*GZMB, GNLY*). A third non-cycling αβ T cell population not expressing *CCL5* was also identified in spleen, termed *CCL5*-αβ T cells. Compared to *CCL5*+ counterparts (inferred to be effector cells) in spleen or bone marrow, *CCL5*-αβ T cells in spleen and *CCL5*-T/ILCs in bone marrow each had higher expression of genes associated with migratory circulation patterns characteristic of naïve and central memory T cells, including expression of lymph node homing gene, *CCR7,* and lymph node egress gene, *S1PR1* [5, 14]. CD4 and CD8 αβ T cell subsets were not differentiated in bone marrow due to the low number of total αβ T cells identified, and gene expression of *CD4, CD8A,* and *CD8B* expression was too sparse in the spleen dataset to accurately differentiate the two αβ T cell subsets.

Though the majority of αβ T cells and ILCs were *CCL5*-in thymus and lymph node, small populations of combined T cells and ILCs that were *CCL5*+ were identified, termed *CCL5*+ T/ILCs in each of thymus and lymph node. Remaining non-cycling *CCL5*-αβ T cell populations in thymus and lymph node were also segregated into CD4 (expressing *CD4*) and CD8 (expressing *CD8A* and *CD8B*) αβ T cell subsets. Within CD4 and CD8 αβ T cell subsets, further differentiation was performed based on expression of the migratory regulator gene, *KLF2*. In both thymus and lymph node, *KLF2*+ CD4 and CD8 αβ T cell populations were presumed to be able to egress from tissues and circulate, often in conjunction with markers characteristic of naïve or central memory T cells, including *IL7R* and *CCR7* in both tissues, as well as *SELL*, *S1PR1,* and *LEF1* in lymph node [5, 14]. In total, a population of *KLF2*+ CD4 αβ T cells and *KLF2*+ CD8 αβ T cells was identified in each of thymus and lymph node, totaling four *KLF2*+ αβ T cell populations identified. A total of seven *KLF2*-αβ T cell populations were identified in thymus and lymph node. In lymph node, a population of *KLF2*-CD8 αβ T cells was identified along with two populations of CD4 αβ T cells lacking *KLF2* expression, termed *KLF2*-CD4 αβ T cells and follicular CD4 αβ T cells. Though both *KLF2*-CD4 αβ T cells and follicular CD4 αβ T cells in lymph node expressed genes such as *CXCR5* and *CD40LG* that may be associated with a follicular location, follicular CD4 αβ T cells had higher expression of additional genes indicating follicular T cell signaling, including *PDCD1, CTLA4, ICOS, PRDM1,* and *IL2RA* [14]. In thymus, *KLF2*-αβ T cells included a population of *KLF2*-CD8 αβ T cells, two populations of *KLF2*-CD4 αβ T cells further divided by *CCR9* expression, and a population of αβ T cells with high expression of *CD1D* and *CD1E*. Expression of *CCR9* on *KLF2*-αβ T cells in thymus suggested a role in thymocyte development or gut homing potential [12, 58, 59]. The final population of *KLF2*-αβ T cells in thymus was termed *CD1D*+*CD1E*+ αβ T cells, but such cells could not be easily discerned as CD4 or CD8 αβ T cells. These cells also had high expression of genes such as *CD1D, CD1E*, *CCR4, GATA3* that were not typically expressed at high levels by other αβ T cell populations.

Similar to αβ T cells, populations comprised solely of ILCs could be divided by *CCL5* expression. Two *CCL5*+ ILC populations were identified in spleen, termed cytotoxic ILCs and *NCR1*+*EOMES*+ ILCs. Cytotoxic ILCs in spleen expressed cytotoxicity-related genes *GZMB* and *GNLY*. Conversely, *NCR1+EOMES+* ILCs in spleen lacked expression of cytotoxicity genes and instead had strong expression of genes characteristic of porcine natural killer (NK) cells [5, 14], including *NCR1*, *EOMES*, and *KLRB1*. Lastly, two populations of *CCL5-* ILCs, identified in each of bone marrow and lymph node, had high expression of *KIT, IL7R,* and *CD69* and were termed *KIT*+ ILCs. *KIT+* ILCs in lymph node also expressed *RORC* and *KLRB1*, suggesting a population of group 3 ILCs similar to those identified in porcine intestine that also have similar expression patterns and represent a population of putative lymphoid tissue inducer (LTi) cells [13, 14].

### 3.2 Two independent porcine thymus scRNA-seq datasets yield similar cell type identities

One recent publication [12] defines porcine thymic cell populations with a high level of granularity attributed to defining immune cell populations. We compared the manual annotation of our thymic dataset to that of previous work [12] to establish a general consensus of cell types annotated in porcine thymus across different published datasets. Comparisons were performed using reference-based mapping and cell label predictions (see methods), where the previously published work was treated as a reference dataset (**Figure 3A**), and our thymic dataset was treated as a query dataset (**Figure 3B**) projected onto the reference.

**Figure 3.**
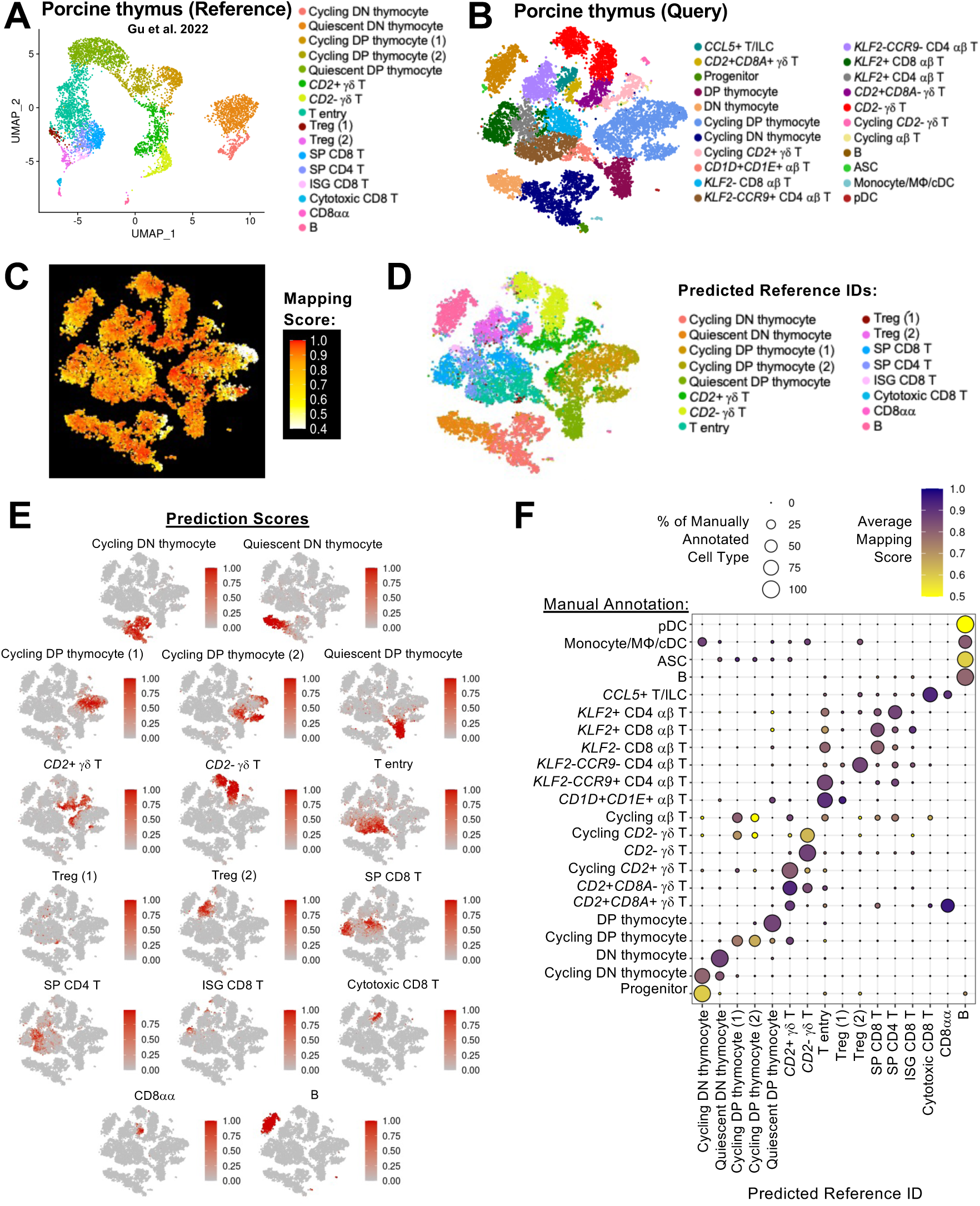
Annotation consensus of porcine thymic cells across scRNA-seq datasets. (A-B) UMAP plot of cell types identified in a previously published dataset of porcine thymus, treated as a reference dataset (A), and t-SNE plot of cell types identified in porcine thymus of this work, treated as a query dataset (B). Each individual point represents one cell, and the color of each point corresponds to annotated cell identity. (C) t-SNE plot of mapping scores (range 0 to 1) obtained from projecting query data in B onto the reference dataset in A. Each individual point represents one cell of the query dataset, and the color of each point corresponds to mapping score, where a higher mapping score indicates better representation of a query cell in the reference dataset. (D) t-SNE plot of predicted reference cell identity obtained from projecting query data in B onto the reference dataset in A. Predicted annotation was assigned as the reference cell type with the highest prediction score. Each individual point represents one cell of the query dataset, and the color of each point corresponds to the predicted reference annotation. (E) t-SNE plots of prediction scores (range 0 to 1) obtained from projecting query data in B onto the reference dataset in A. Each individual point represents one cell of the query dataset, and the color of each point corresponds to the prediction score for cell type annotations found in the reference dataset. (F) Dot plot showing predicted reference cell type (x-axis) for cell types of query data (y-axis). Predicted annotation was assigned as the reference cell type with the highest prediction score. Dot size indicates the percentage of a query cell type (y-axis) predicted to be a reference cell type (x-axis). Dot color indicates average mapping score for those cells predicted as a reference cell type within an annotated query cell type. A higher mapping score indicates better representation of query cells in the reference dataset. Abbreviations: ASC (antibody-secreting cell); cDC (conventional dendritic cell); DN (double negative); DP (double positive); ILC (innate lymphoid cell); ISG (interferon-stimulated gene); MΦ (macrophage); pDC (plasmacytoid denritic cell); scRNA-seq (single-cell RNA sequencing); SP (single positive); t-SNE (t-distributed stochastic neighbor embedding); Treg (T regulatory); UMAP (uniform manifold approximation and projection)

Mapping scores were recovered as a metric for each query cell (**Figure 3C**) and indicated the degree of representation each query cell had in the reference dataset, with higher mapping scores indicating better representation. Predicted reference dataset identities for each cell of the query data were also calculated (**Figure 3D**) by identifying the reference identity with the highest prediction score (**Figure 3E**) for each query cell. Results indicated a general consensus of annotation cell groupings between reference and query datasets with some exceptions (**Figure 3F**). General annotation consensus was found across datasets for thymocytes, gd T cells, mature ab T cell populations, and B lineage cells. Largest discrepancies arose for query populations of progenitor cells, monocyte/MΦ/cDCs, and pDCs, as similarly annotated populations were not identified in the reference data. Query progenitor cells were most closely predicted as DN thymocytes, though with low mapping scores indicating a similar cell type was not represented in the reference data. Monocyte/MΦ/cDCs and pDCs in the query datasets were predicted as most similar to reference B cells, though mapping scores were particularly low for pDCs, again indicating lack of a similar cell type in the reference data. Another interesting finding was *CD1D+CD1E*+ αβ T and *KLF2-CCR9*+ CD4 αβ T cells in the query data being predicted as T entry cells in the reference. High mapping scores were also recovered for the two populations highly predicted as T entry cells, indicating representation of very similar cell populations across both datasets. Results thus indicated a general consensus for most cell types similarly annotated across the two datasets, though populations such as progenitors, monocyte/ MΦ/cDCs, and pDCs may be unique to our newly-established thymus scRNA-seq data, which contains a greater number of cells than previously-published work. Comparison to the previously-published thymus data also suggested populations of T entry cells were differentiated into two distinct populations in our work, including *CD1D+CD1E*+ αβ T and *KLF2-CCR9*+ CD4 αβ T cells.

### 3.3 Two populations of splenic innate lymphoid cells are transcriptionally similar in pigs and humans

Reference-based mapping and prediction is also a useful tool for comparative immunology, where cell types can be compared across single-cell datasets of different species, including between pigs and humans [5, 14, 18, 60, 61]. To perform comparative analysis of splenic cells from pigs and humans, our spleen dataset was mapped to a previously-published scRNA-seq dataset of human spleen [26]. Comparisons were performed using reference-based mapping and cell label prediction (see methods), where the previously published work was treated as a reference dataset (**Figure 4A**), and our splenic dataset was treated as a query dataset (**Figure 4B**) projected onto the reference. Mapping scores were generally high for cell types annotated in our pig query data (**Figure 4C**), indicating pig cells had transcriptionally-similar human counterparts for most splenic cell types, including ILCs. ILCs are not as well-studied in pigs as other species such as humans or mice [14, 62–65], and we characterized two newly-defined splenic populations of ILCs in porcine spleen (**Figure 1D & Figure 2C**), including splenic cytotoxic ILCs and *NCR1+EOMES+* ILCs. Prediction of porcine splenic ILCs to human splenic cell annotations indicated porcine cytotoxic ILCs were primarily predicted as human *FCGR3A*+ NK cells, and porcine *NCR1*+*EOMES*+ ILCs were primarily predicted as *CD160*+ NK cells (**Figure 4D-E**), with high mapping scores indicating representation of highly similar ILCs and NK cells across datasets (**Figure 4F**). In a new dataset, porcine and human ILC/NK cells were subsetted and integrated, revealing hierarchical clustering of annotated cells resulted in species intermixing, with one node containing porcine cytotoxic ILCs and human *FCGR3A*+ NK cells that were most closely related to one another, while the other node included porcine *NCR1+EOMES*+ ILCs and human *CD160*+ NK cells that were most closely related to one another (**Figure 4G**). Visualization with dimensionality reduction supported these relationships, again indicating pig-to-human similarities of porcine cytotoxic ILCs to human *FCGR3A*+ NK cells and porcine *NCR1+EOMES*+ ILCs to human *CD160*+ NK cells (**Figure 4H**). Collectively, comparison of pig and human ILC/NK cell populations revealed conservation across species, with two primary populations identified in spleen belonging to the ILC lineage that encompasses both ILC subsets and NK cells [66, 67]. Porcine splenic ILCs did not meet the traditional classification of porcine NK cells due to lack of specific markers such as *CD8A* but did have gene expression best represented by the group 1 ILC lineage, reminiscent of previous work in porcine ileum [14]. Specific classification of species-conserved populations of ILCs remains complicated due to tissue-specific features, as well as transcriptional and functional overlapping of subsets, especially between ILC1s and NK cells [66–68]. However, cross-species comparisons reveal high levels of transcriptional conservation for splenic ILCs in pigs and humans regardless of ILC1 versus NK classification.

**Figure 4.**
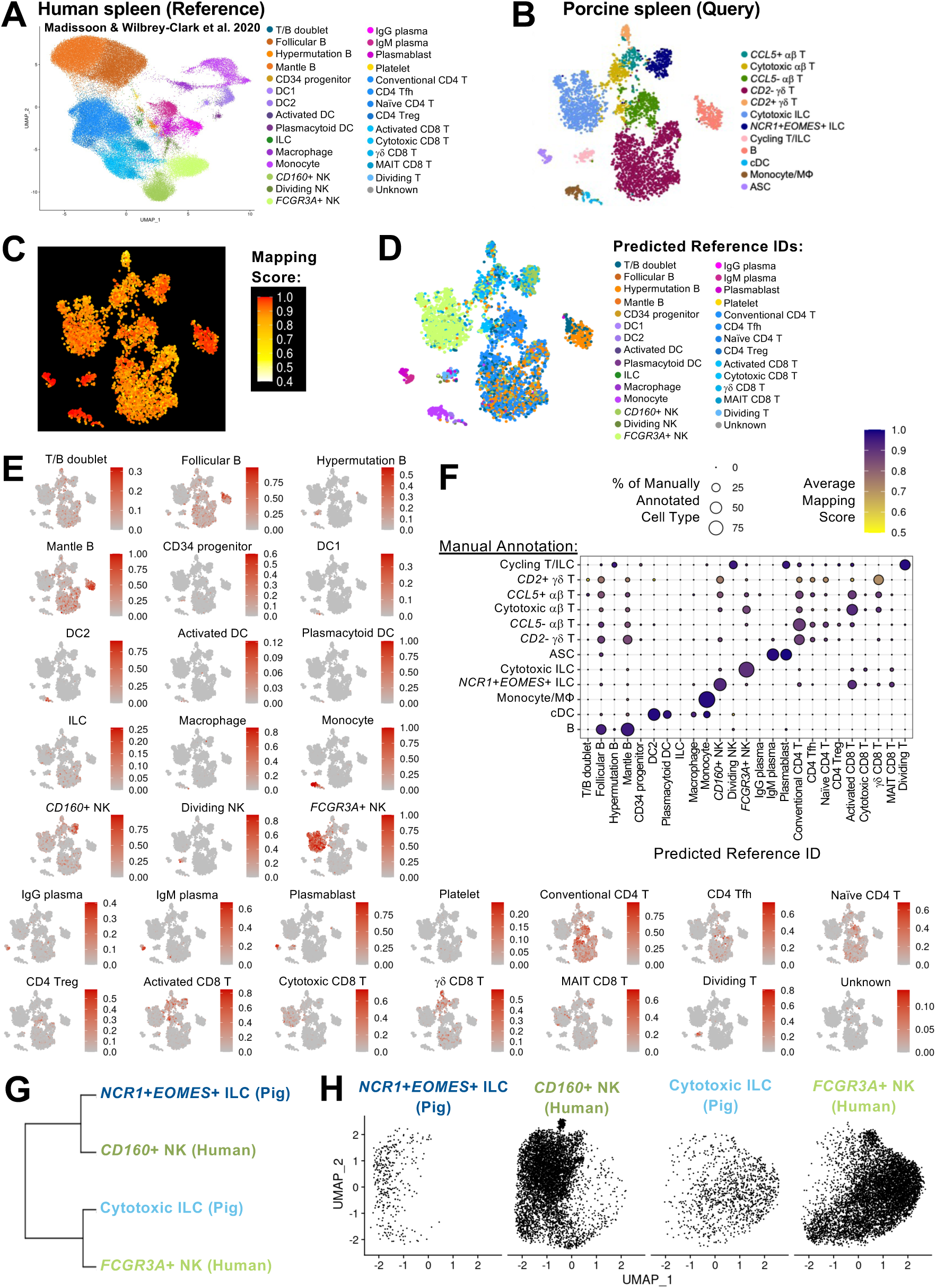
Porcine splenic ILCs share transcriptional similarities with NK cells in human spleen. (A-B) UMAP plot of cell types identified in a previously published scRNA-seq dataset of human spleen, treated as a reference dataset (A), and t-SNE plot of cell types identified in porcine spleen of this work, treated as a query dataset (B). Each individual point represents one cell, and the color of each point corresponds to annotated cell identity. (C) t-SNE plot of mapping scores (range 0 to 1) obtained from projecting query data in B onto the reference dataset in A. Each individual point represents one cell of the query dataset, and the color of each point corresponds to mapping score, where a higher mapping score indicates better representation of a query cell in the reference dataset. (D) t-SNE plot of predicted reference cell identity obtained from projecting query data in B onto the reference dataset in A. Predicted annotation was assigned as the reference cell type with the highest prediction score. Each individual point represents one cell of the query dataset, and the color of each point corresponds to the predicted reference annotation. (E) t-SNE plots of prediction scores (range 0 to 1) obtained from projecting query data in B onto the reference dataset in A. Each individual point represents one cell of the query dataset, and the color of each point corresponds to the prediction score for cell type annotations found in the reference dataset. (F) Dot plot showing predicted reference cell type (x-axis) for cell types of query data (y-axis). Predicted annotation was assigned as the reference cell type with the highest prediction score. Dot size indicates the percentage of a query cell type (y-axis) predicted to be a reference cell type (x-axis). Dot color indicates average mapping score for those cells predicted as a reference cell type within an annotated query cell type. A higher mapping score indicates better representation of query cells in the reference dataset. (G) Hierarchical clustering of indicated porcine ILC and human NK cell types when established as a merged dataset. (H) UMAP plot of indicated porcine ILC and human NK cell types when established as a merged dataset. Cells for each cell type are shown in independent panels where each point represents one cell. Abbreviations: ASC (antibody-secreting cell); cDC (conventional dendritic cell); DC (dendritic cell); DN (double negative); DP (double positive); ILC (innate lymphoid cell); MΦ (macrophage); MAIT (mucosal-associated invariant T); NK (natural killer); scRNA-seq (single-cell RNA sequencing); t-SNE (t-distributed stochastic neighbor embedding); Tfh (T follicular helper); Treg (T regulatory); UMAP (uniform manifold approximation and projection)

### 3.4 Porcine lymph nodes share structural and signaling similarities to humans

Lymph nodes are critical tissue sites for immune induction where immune responses are dependent on the intricate organization of follicles that foster cellular interactions required for immune processes. To better understand cellular organization in the porcine lymph node, communication networks were calculated for annotated cell types of the lymph node scRNA-seq dataset and used to construct an inferred three-dimensional organization of cells based on expression of genes encoding ligands and receptors of known signaling pathways (**Figure 5A**). Based on reconstructed cellular organization patterns, significant inferred interactions and non-interactions were found for all cell-cell combinations (**Figure 5B**), and the distances of each cell type from the center of the inferred three-dimensional structure were also calculated (**Figure 5C**). The three-dimensional structure resembled a follicle, where cell density was highest at the structure center and contained the largest numbers of B cells (**Figure 5D**). Proximity of *KLF2*-T cell and ILC populations in the B cell-dense center further suggested similarities to follicular organization. Loss of *KLF2* expression is associated with T cell activation, tissue retention, and lymph node follicle entry [69], and lymph node T cell and ILC populations with low/no expression of *KLF2* (*KLF2-* CD8 αβ T, *KLF2-* CD4 αβ T, follicular CD4 αβ T, *KIT*+ ILC*, CCL5*+ T/ILC; **Figure 2C**) generally had increased numbers of significant interactions with other cell types (**Figure 5B**) and were located in greatest numbers closer to the structure center (**Figure 5C-D**), suggesting cells with little to no *KLF2* expression were follicle-associated and may have important roles in germinal center signaling interactions with other cell types. Several of the strongest recovered cell-cell signaling pathways (CD45, MHC-II, CD40, CD86, CD80, PDL2, PD-L1) are involved with germinal center immune processes [70–74] (**Figure 5E**). Monocyte/MΦ/cDCs alongside subsets of CD4 αβ T cells and B cells were identified as particularly important senders (expressing ligand) or receivers (expressing receptor) of signaling pathways associated with germinal center processes (**Figure 5F**). These three general cell groupings (monocytes/MΦ/cDCs, CD4 αβ T cells, and B cells) are also those traditionally defined to interact via ligand-receptor bindings to activate an antigen-specific adaptive immune response in lymph node germinal centers [70–74].

**Figure 5.**
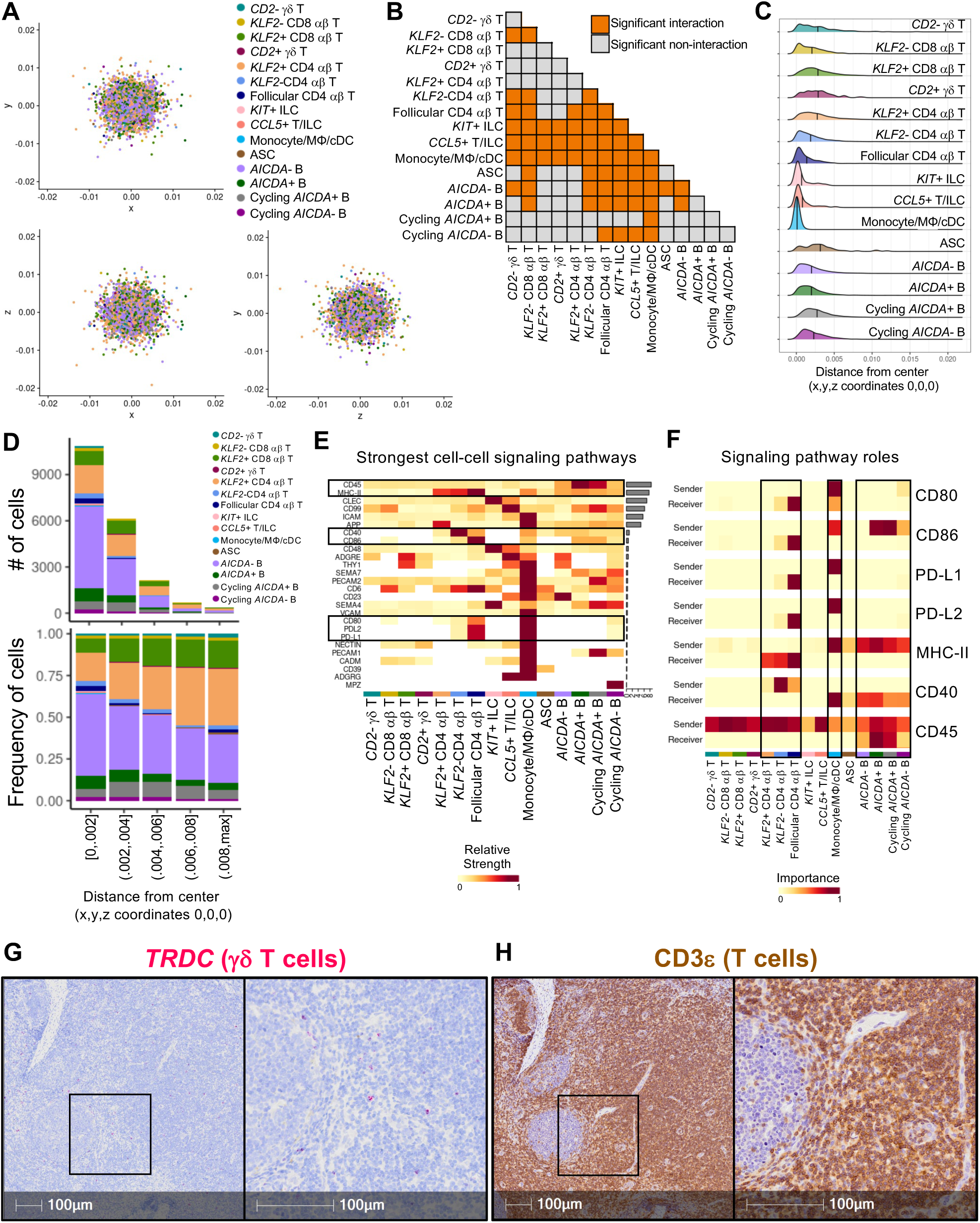
Reconstruction of lymph node signaling and three-dimensional organization indicates porcine lymph nodes undertake conventional germinal center immune induction signaling processes with minimal contributions from γδ T cells. (A) Three-dimensional spatial coordinates (x,y,z axes) of lymph node cellular organization inferred from expression of ligand- and receptor-encoding signaling genes expressed by individual cells of the pig lymph node scRNA-seq dataset. Each individual point represents one cell, and the color of each point corresponds to annotated cell identity. (B) Table of inferred significant interaction (orange) or non-interaction (grey) between lymph node cell types. Significant interaction or non-interaction were inferred based on significant high or low proximity, respectively, of cells in the three-dimensional structure shown in (A). No non-significant interactions were noted for any combination of annotated cell types. (C) Ridge plot showing the distribution of cell distances from the central coordinates (0,0,0) of the three-dimensional structure in (A). Distributions are plotted for each individual cell type, and mean distance from central coordinates is noted with a vertical line. (D) Stacked bar plots showing the number of cells (top) or frequency of cells (bottom) (y-axes) occurring at different ranges of distance from the central 0,0,0 coordinates of the structure shown in (A) (x-axes). Bar color corresponds to an annotated cell type. (E) Heatmap of the strongest cell-cell signaling pathways (y-axis) inferred amongst cell types (x-axis) in porcine lymph node scRNA-seq data. Fill color corresponds to the relative strength a corresponding cell type has in a corresponding signaling pathway. To the right of the heatmap, bar plots indicate the overall strength of each signaling pathway. (F) Heatmaps indicating roles of annotated lymph node cell types (x-axes) as senders (expressing ligand-encoding genes) or receivers (expressing receptor-encoding genes) of signals (y-axes) for a subset of signaling pathways taken from (E). Fill color indicated the importance of a corresponding cell type in acting as a sender or receiver for an indicated pathway. (G-H) Microscopy images of *TRDC* RNA staining (red) to indicate presence of γδ T cells (G) and CD3ε protein staining (brown) to indicate presence of total T cells (H) in a section of lymph node also taken and processed for scRNA-seq. Abbreviations: ASC (antibody-secreting cell); cDC (conventional dendritic cell); ILC (innate lymphoid cell); MΦ (macrophage)

Within CD4 αβ T cell subsets, follicular CD4 αβ T cells were generally the most important recipients of signals from several pathways, including CD80, CD86, PD-L1, PD-L2, and MHC-II signaling followed by *KLF2*-CD4 αβ T cells, while *KLF2+* CD4 αβ T cells had little to no role as receivers for most pathway signals (**Figure 5F**). Greater importance of follicular CD4 αβ T cells, followed by *KLF2*-CD4 αβ T cells, in several germinal center signaling processes (**Figure 5F**) coincided with increased numbers of significant interactions with other cell types (**Figure 5B**) and increased association with structure center where other follicle-associated cell types were located (**Figure 5C-D**) compared to *KLF2*+ CD4 αβ T cells that had more significant non-interactions (**Figure 5B**) and were more often identified further from the structure center (**Figure 5C-D**). Results strongly suggest the inclusion of cells annotated as follicular CD4 αβ T cells were indeed associated with lymph node follicles and are critical to signaling processes required for germinal center processes. Results also suggest *KLF2*-CD4 αβ T cells have increased associations with lymph node follicles and associated signaling pathways compared to their *KLF2*+ counterparts, suggesting *KLF2* may be a marker that can be targeted for functional inference of porcine CD4 T cells in future work.

Though many of the interactions and signaling pathways defined in porcine lymph node were similar to those defined in other species [70–74], a peculiarity of pigs is an increased number of γδ T cells compared to other species, including in lymph node tissue [75, 76].Annotated γδ T cell populations in our porcine lymph node scRNA-seq data had mostly non-interactions with other cell types (**Figure 5B**), were generally located outside of the reconstructed structure center (similar to other *KLF2*+ T/ILC populations; **Figure 5C-D**), and were negligible contributors to most lymph node cell-cell signaling pathways (**Figure 5E-F**). *In situ* staining for γδ T cell marker, *TRDC,* verified localization of γδ T cells outside of germinal centers (**Figure 5G**), and γδ T cells made up a minor fraction of total T cells (indicated by CD3ε immunolabeling) in lymph node tissue (**Figure 5H**) [75, 77]. Collectively, *in situ* staining and scRNA-seq analysis results suggested γδ T cells are minimal contributors to germinal center immune induction in pig lymph nodes. Thus, though increased percentages of γδ T cells in pig lymph nodes presents a peculiarity dissimilar to human lymph nodes, histological staining and *in silico* reconstruction of lymph node structure and signaling networks suggest porcine γδ T cells have little influence on fundamental germinal center processes.

### 3.5 Using an open-source application interface of the porcine single-cell immune atlas to explore immune cells in pig bone marrow

To promote usability of the multi-tissue immune atlas created in this work, we developed an open-source application interface available to perform gene expression query and cell type population comparisons across pig primary and secondary lymphoid organs captured in our work. The web application, called Shiny-PIGGI will be an important tool for exploration of porcine immune genes and cell types in these tissues.

As a demonstration, we utilize the application to interrogate immune cell types recovered from bone marrow and to compare them to cells in other tissues of our immune atlas. In **Figure 6A**, we show the home page for the Shiny-PIGGI web application, along with key functionalities of each tab, including differential gene expression, gene expression query, reference cell type mapping. The homepage also includes a link to downloadable data files for easy downstream analysis and visualization and a “Differential Gene Expression” section for performing user-specified cell type-specific differential gene expression analysis.

**Figure 6.**
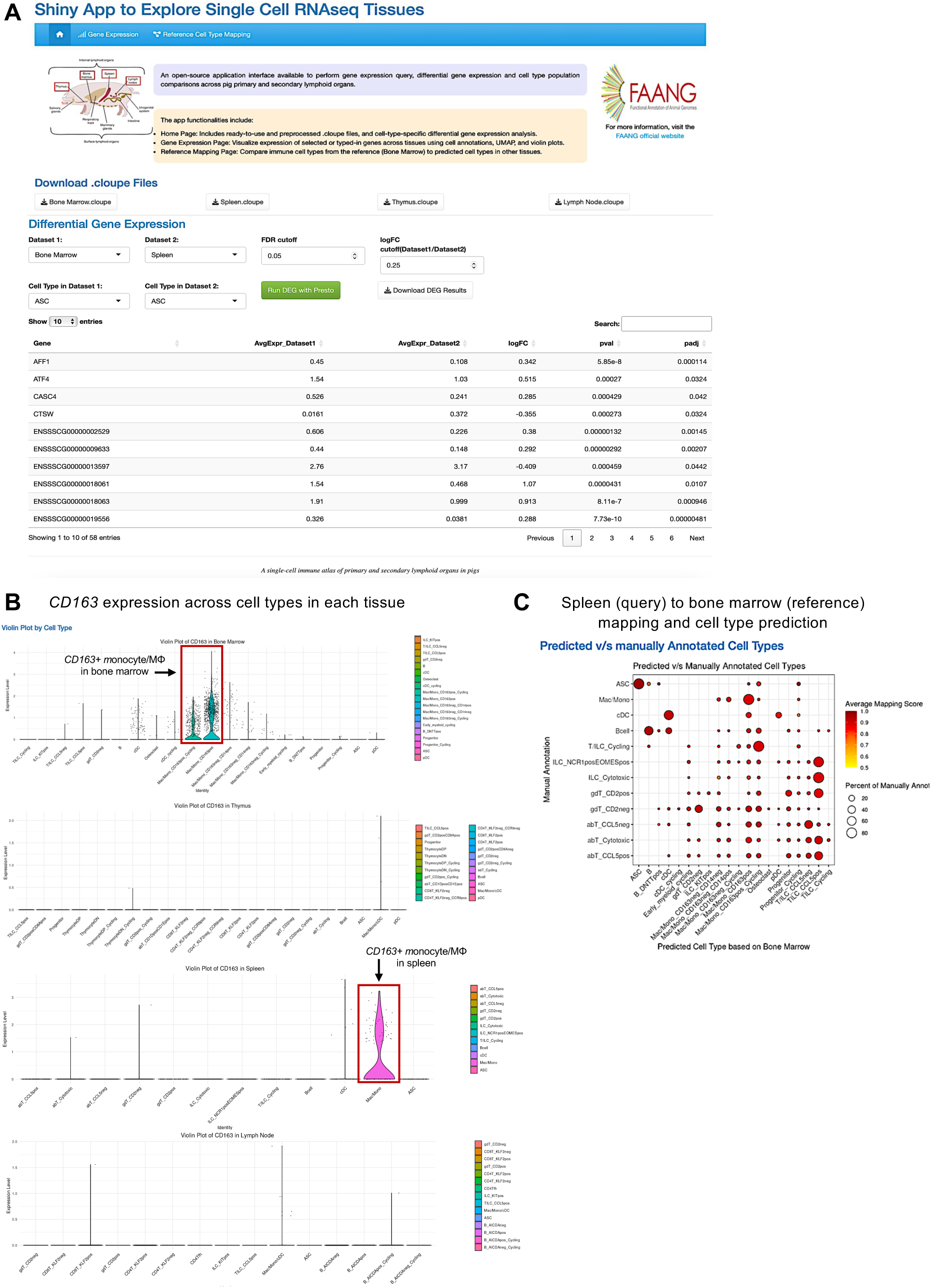
Utilization of an interactive application for further data query of monocyte/macrophage heterogeneity in pig bone marrow. (A) Home page for the web application created to query the pig single-cell multi-organ immune atlas. (B) Violin plots created to assess *CD163* expression in cell types of all tissues under the “Gene Expression” tab of the web application. (C) Dot plot showing predicted bone marrow reference cell type (x-axis) for cell types of spleen query data (y-axis) as created under the “Reference Cell Type Mapping” tab of the web application. Predicted annotation was assigned as the reference cell type with the highest prediction score. Dot size indicates the percentage of a query cell type (y-axis) predicted to be a reference cell type (x-axis). Dot color indicates average mapping score for those cells predicted as a reference cell type within an annotated query cell type. A higher mapping score indicates better representation of query cells in the reference dataset.

Under the “Gene Expression” tab, we demonstrate how the application interface can be used to assess expression of user-specified genes across all four dataset tissues, such as key marker gene *CD163* that was used to annotate unique monocyte/macrophage populations in bone marrow. Violin plots of *CD163* expression within cell types of each tissue that were generated in the web application are shown in **Figure 6B** and indicate *CD163* was not highly expressed by myeloid populations in thymus or lymph node; however, cycling and non-cycling *CD163*+ monocyte/MΦ populations were recovered in bone marrow, and some splenic monocytes/MΦs also expressed *CD163*. Further comparison of cell types recovered from bone marrow and spleen datasets was performed using mapping procedures similar to those used in **Figures 3 & 4** under the “Reference Cell Type Mapping” tab in the web application, where bone marrow was considered as a reference dataset, and spleen was considered as a query dataset. **Figure 6C** shows a dot plot obtained from the application interface results of spleen-to-bone marrow reference mapping. Splenic cells had generally high mapping scores to the bone marrow dataset, indicating close counterparts could be identified in bone marrow. Similarly annotated cell types were generally in consensus based on cell type predictions, including B cells, ASCs, monocytes/MΦs, and cDCs. Remaining splenic cell populations belonged to the T/ILC lineage and were also largely predicted as types of T/ILCs in bone marrow data; however, cycling T/ILCs in spleen were predicted as mostly cycling *CD163*+ monocytes/MΦs from bone marrow, likely due to shared expression of many cell cycle genes that drive transcriptional profiles in both populations. Other effector populations of T cells and ILCs described in spleen were predicted mostly as *CCL5*+ T/ILCs from bone marrow, coinciding with *CCL5+* T/ILCs being a likely mature effector cell population in bone marrow. Overall, the application is an open-source resource that can be used to query tissue datasets for forming important biological conclusions and inferences, such as examples shown above. Access to the query interface is described in the data availability statement of this work.

## 4 Discussion

Results described above demonstrate the vast heterogeneity of the immune landscape across primary and secondary lymphoid organs in pigs and include important insights into cell phenotypes, functions, and implications for comparisons across species. Thymus and bone marrow, the two primary lymphoid organs analyzed, were home to progenitor cells, cells undergoing recent lineage commitment and development, as well as fully differentiated cells and potential tissue-resident cell populations. Comparison of thymus cells to another porcine thymus scRNA-seq dataset revealed further functional implications for several cell subsets, including high degrees of validatory, functionally-implicating annotations across datasets along with novel cell populations captured in our larger thymus scRNA-seq dataset. In bone marrow, we demonstrate similarities and differences to cells captured at other tissue sites through interactive query with a newly developed application interface. In the two secondary lymphoid organs analyzed, spleen and lymph node, diverse effector cell subsets were discovered and were highly distinct across the two immune tissues, though cells from both tissues shared similarities with human cells from corresponding lymphoid organs. In spleen, several highly activated, effector T and ILC populations were characterized, including novel ILC subsets highly similar to those in human spleen. In lymph node, conventional leukocyte populations that facilitate germinal center immune reactions were characterized and had highly similar signaling and organizational dynamics to those in human lymph nodes. Overall, our work provides an expanded overview of the immune landscape in pigs across primary and secondary lymphoid organs, and each tissue was critically analyzed in a different manner to demonstrate the utility of the single-cell immune atlas as a resource that can be wielded to address further research queries through a variety of methodologies. Furthermore, an interactive web application for data query was created and is publicly available to enable better accessibility and usage of our dataset as a resource, including for non-computational viewership and query.

Though our work successfully captures and analyzes a diverse array of immune cells across multiple immunologically-critical anatomical locations of pigs, the atlas is by no means comprehensive. Cells were captured from two animals and are not reflective of the full spectrum of pig life stages and biological scenarios, including age, rearing environment, disease state, gender, or genetic predispositions. Some cell types were captured at low abundances and other populations may have potentially been excluded due to rarity or biased capture from cell isolation processes used (e.g. mechanical rather than enzymatic digestion, cryopreservation, or on-column dead cell removal methods that were utilized in our samples). Lastly, additional anatomical locations likely include expanded immune cell diversity, such as mucosal effector sites lacking organized primary or secondary lymphoid structures (e.g. lungs, intestinal tract), additional lymph nodes draining unique effector sites, or even neurological tissues. Regardless, our work presents an immunologically-focused multi-tissue, single-cell atlas in pigs and serves as an initial data resource to expand upon for understanding the comprehensiveness of the porcine immune system. Further, this work expands the number of tissues interrogated through single-cell analyses for the pig, increasing the value of large scale integration of such data for accurate cell annotations and foundation models of biology at the cell level [78].

In that vein, this work was completed as part of the Functional Annotation of Animal Genomes (FAANG) research initiative to annotate the functional components of domesticated species through detecting RNA expression and epigenetic status in tissues [79]. Describing the transcriptome of important cell types within tissues and under conditions relevant to understanding immune response and function is a first step to develop immune system gene regulatory knowledge, which has been initiated in this work. The second step is understanding the interactions of components of the system by linking genes with their regulators through these regulatory elements; a network describing such interactions as an “intermediate phenotype” is a paradigm for understanding complex disease and predicting biological outcomes [80, 81]. Global transcriptomic studies in FAANG research are expected to reveal co-expression of transcriptional regulators with their target genes in specific tissues or cells [82], providing useful annotation of livestock genomes, and this work may be further applied in future as a resource to reach such goals. The datasets generated herein may also serve as resources for various other research queries such as those related to pig immunology and health or potential biomedical applications of pigs for human health advancement.

## 6 Author Contributions

**Jayne E. Wiarda:** conceptualization, data curation, formal analysis, investigation, methodology, project administration, resources, software, visualization, writing – original draft, writing – review & editing. **Muskan Kapoor:** data curation, methodology, software, visualization, writing – original draft, writing – review & editing. **Sathesh K. Sivasankaran:** data curation, software, writing – review & editing. **Kristen A. Byrne:** investigation, methodology, writing – review & editing. **Crystal L. Loving:** conceptualization, data curation, investigation, methodology, project administration, resources, supervision, writing – original draft, writing – review & editing. **Christopher K. Tuggle:** conceptualization, investigation, funding acquisition, project administration, resources, supervision, writing – original draft, writing – review & editing.

## 8 Acknowledgments

We thank (1) Dr. Juber Herrera-Uribe and Zahra Bond for sample preparation support; (2) Drs. David Alt and Mike Baker for sequencing services; (3) Dr. Darrell Bayles for data transfer and management; (4) Drs. Lance Daharsh and Daniel Nielson for technical input; (5) Adrienne Shircliff and Judith Stasko for histology services; (6) Alexey Sokolov at EMBL-European Bioinformatics Institute for help in submitting data reported in this manuscript to the European Nucleotide Archive.

## 9 Data availability statement

Scripts used for data analysis are available at https://github.com/SwiVi/FAANG_MultiTissue_Immune_scRNAseq. The interactive query application, Shiny-PIGGI, can be accessed at https://shinypiggi.ansci.iastate.edu/ or downloaded from https://github.com/kapoormuskan/Pig_Immune_Tissue_ShinyApp. Raw sequencing data are available at the European Nucleotide Archive (ENA) under project PRJEB97326 (https://www.ebi.ac.uk/ena/browser/view/PRJEB97326). Additional files of processed datasets used for computational analysis and files compatible with interactive data query through Loupe Browser (10X Genomics; .cloupe files) are available at Ag Data Commons (https://www.doi.org/10.15482/USDA.ADC/29492726.v1).

